# An artistic approach to neurofeedback for emotion regulation

**DOI:** 10.1101/794560

**Authors:** Damien Gabriel, Thibault Chabin, Coralie Joucla, Thomas Bussière, Aleksandra Tarka, Nathan Galmes, Alexandre Comte, Guillaume Bertrand, Julie Giustiniani, Emmanuel Haffen

## Abstract

Neurofeedback has been shown to be a promising tool for learning to regulate one’s own emotions in healthy populations and in neuropsychiatric disorders. While it has been suggested that neurofeedback performance improves when sensory feedback is related to the pathology under consideration, it is still difficult to represent in real time a proper feedback representative of our emotional state. Since emotion is a central part of people’s dealings with artworks, we have initiated a collaboration between neuroscientists and artists to develop a visual representation of emotions that can be used in neurofeedback experiences. As a result of this collaboration, emotions were represented as particles, moving in a white sphere according to valence and arousal levels. In this study, several possibilities for particle control were explored: direction of particles, their concentration in a specific place, or their gravity. 107 participants evaluated these performances, either in laboratory condition or at various scientific and artistic events. At the end of the experiment, questionnaires were distributed to participants who were asked to indicate on scales ranging from 0 to 5 how artistic the different representations were and could be used as a clinical activity, whether they thought they had successfully controlled the particles during the neurofeedback exercise, and whether they had appreciated the experience. We found that influing on the direction and concentration of particles was considered the most artistic with an average score around 3/5. 47% of the participants considered the concentration of particles as artistic. In addition, although this is not the purpose of this study, we found that participants could significantly control the direction of particles during this session. These encouraging results constitute a first step before evaluating the effectiveness of our emotional neurofeedback over several sessions in healthy, then pathological populations.

## 1. Introduction

Neurofeedback is a technique that consists in measuring in real time a neurophysiological activity in order to extract a parameter of interest and present it to the participant, typically via visual or auditory feedback. The purpose is to teach the participant to modify this parameter. Neurofeedback can be used to improve cognitive performance, such as memory, attention or emotions (Gruzelier 2014; Gaume et al. 2016). It can be used in healthy people (e.g. (Gruzelier 2018)) but it is mainly perceived as a therapeutic tool for the treatment of mental disorders (e.g. epilepsy, attention disorders, addiction, depression). There are at least two ways in which regulating brain activity by neurofeedback can be beneficial for the treatment of mental disorders. Self-regulatory training can focus on an abnormal process, such as hyper- or hypoactivation of specific brain areas or networks. But neuromodulation can also act in another way, by activating or suppressing circuits that do not function abnormally, but whose neuromodulation can nevertheless produce clinical benefits (D. Linden 2013). This implies that clinical benefits can be achieved through self-regulatory training that activates compensatory circuits or inhibits circuits that appear normal when viewed in isolation but contribute to pathology-related dysfunction (D. E. J. Linden 2014).

The important parameters to consider when conducting a neurofeedback experiment are the method of measuring neurophysiological activity, the brain areas to be targeted, and the type of feedback to be presented to participants. All these parameters obviously depend on the phenomenon we want to study. The two main techniques for measuring neurophysiological activity in neurofeedback are electroencephalography (EEG) and functional magnetic resonance imaging (fMRI). EEG neurofeedback consists in measuring the power of cerebral electrical activity at frequency bands of interest on a few electrodes placed on the surface of the scalp, with a time accuracy of the order of a millisecond. EEG neurofeedback has the advantage of being easy to use and can be performed ambulatory. Neurofeedback by fMRI is a relatively recent development of neurofeedback based on blood oxygenation contrasts from the Blood-Oxygen-Level-Dependent (BOLD) signal (for reviews, see (deCharms 2008; Sulzer et al. 2013; Weiskopf 2012)). Neurofeedback training by fMRI can overcome some of the limitations of traditional forms of neurofeedback in EEG, thanks to its higher spatial resolution and the integration of the entire brain. This approach is non-invasive, spatially accurate, and capable of targeting deep brain structures such as the amygdala. Unlike EEG neurofeedback, the fMRI technique does not really provide ‘real-time’ feedback because of the hemodynamic delay of about 5 seconds between current neural activity and the vascular response that creates the fMRI signal. However, this delay is not an obstacle to neurofeedback when participants receive this information prior to the experiment (Weiskopf et al. 2004; D. E. J. Linden 2014).

The cerebral area to measure and to be controlled by the participant is a parameter that depends on the phenomenon to be studied and is defined from the existing literature in the field. In the case of using neurofeedback to learn how to regulate emotions, most EEG neurofeedback studies focus on the activity of the prefrontal cortex, which acts as a modulator of primary emotional responses, through its connections with deep brain structures (Spielberg et al. 2012). Dominant activity in right versus left prefrontal areas is associated with withdrawal behavior and negative emotions, while opposite representation (i.e., higher activity on the left versus right) accompanies approach behaviors and positive emotions (Davidson 1988; 1998; Papousek et al. 2014). Thus, the alpha frontal asymmetry recorded in the EEG reflects functional differences between approach and avoidance motivation systems (see as reviews (Coan et Allen 2004; Davidson 1998; 1992; Harmon-Jones et Gable 2018; Sutton et Davidson 1997)). Since alpha power is assumed to reflect a decrease in metabolic activity (Cook et al. 1998; Davidson et al. 1990), reduced alpha activity in right prefrontal electrodes is associated with negative emotions, for example after viewing unpleasant films (Papousek et al. 2014; Wheeler, Davidson, et Tomarken 1993). On the other hand, reduced alpha activity on the left is related to positive emotions, for example after viewing happy movies or listening to pleasant music (Wheeler, Davidson, et Tomarken 1993; Arjmand et al. 2017). Several case studies have shown the effectiveness of training to control alpha asymmetry to reduce depressive symptoms (Baehr et Baehr 1997; Baehr, Rosenfeld, et Baehr 1997; Choi et al. 2011; Peeters et al. 2014). Frontal asymmetries associated with emotions and motivation have also been observed at the Theta band level (e.g. (Aftanas et Golocheikine 2001; Ertl et al. 2013)) and at the upper beta band level (e.g. (Paquette, Beauregard, et Beaulieu-Prévost 2009; Pizzagalli et al. 2002). In addition to the measurement of emotional valence, Ramirez and Vamvakousis added an additional parameter in calculating the emotional arousal, in order to conform to Russell’s emotional representation model (Ramirez et Vamvakousis 2012). Arousal is calculated as the ratio between beta and alpha bands at the prefrontal cortex, and when associated with valence, it offers the possibility to have a bidimensional representation of emotions. In fMRI, neurofeedback techniques target deep brain structures that cannot be recorded in the EEG, such as the amygdala or the insula, which play a major role in motivational approach and avoidance systems (e.g. (Cunningham, Raye, et Johnson 2005; Cunningham et al. 2010; Schlund et Cataldo 2010; Spielberg et al. 2012)). Several pilot studies have explored the feasibility of training to regulate emotions with fMRI neurofeedback in patients with neuropsychiatric disorders. These studies focused on the self-regulation of the anterior insula (Caria et al. 2007; 2010) in schizophrenic patients (Ruiz et al. 2013), and the self-regulation of the left amygdala (Zotev et al. 2011; 2013) in patients with bipolar or depressive disorders (Young et al. 2014). While training to over-regulate amygdala activity had a potentially positive effect on depressed patients (Young et al. 2014; Yuan et al. 2014), training to under-regulate may help reduce amygdala hyperactivation and improve emotional regulation in patients with bipolar disorder. The combination of simultaneous recordings in EEG and fMRI in the self-regulation of emotions has also been explored (Cavazza et al. 2014; Kinreich et al. 2014; Meir-Hasson et al. 2014; Shtark et al. 2015; Zich et al. 2015). Cavazza and colleagues found an increase in BOLD activity in the prefrontal cortex while subjects regulated their frontal asymmetry in neurofeedback (Cavazza et al. 2014). Similarly, a correlation between the laterality of the BOLD signal at the amygdala and the level of alpha-frontal asymmetry has been observed (Zotev et al. 2016).

The last parameter to be taken into account, namely the sensory feedback presented to the participants, is still little explored. Remarkably, psycho-sociological factors, particularly motivational factors, which also have a major influence on the potential clinical effectiveness of neurofeedback, have been poorly evaluated. Thus, whatever the pathology considered, the majority of neurofeedback tasks are tedious, with brain activity frequently represented in the form of histograms whose level rises or falls in real time. More playful neurofeedback applications, such as video games, have also been developed, but are not related to the pathology to treat, which raises questions about their effectiveness. It has already been pointed out that traditional approaches to brain studies do not take into account the specificities of each individual (Bagdasaryan et Quyen 2013). Thus, it is likely that a Neurofeedback approach will have to adapt to the pathology of interest. Exploratory approaches to representing feedback in relation to the activity you want to improve have been put in place. For example, using neurofeedback to optimize the performance of actors, participants saw themselves on stage thanks to 3D glasses and the control of their brain activity made possible to vary the brightness of the scene and reduce the noise of the audience (Gruzelier 2014). In the context of emotions and the management of emotional disorders, representing feedback related to the pathology is much more complex because it raises the question about the possibility to represent visually or auditorily an emotion. Since emotion is a central part of people’s dealings with artworks, first approaches have been tested in this direction, for example with color schemes that vary when one must feel tenderness or anxiety (Lorenzetti et al. 2018). Ramirez and colleagues performed a musical neurofeedback task for treating depression in elderly people. In that study, participants could manipulate musical parameters in real time by increasing the volume of music with a high arousal state and increasing the tempo when the valence level also increased (Ramirez et al. 2015).

As part of this project, we have initiated a collaboration between scientists and digital artists to develop a visual representation of emotions that can be used in neurofeedback experiments. For this purpose, it was necessary that, in addition to being artistic, the feedback provided to participants be controllable, and therefore it can be used in a clinical activity. To establish a visual representation of emotions, the artists involved in the project started from the very definition of the word emotion. The term emotion has an active connotation since it derives from the Latin word *emovere*, to set in motion (which gives the terms movement, motivation). Thus, emotions were represented as moving particles slightly tinged according to their location and moving in a white sphere. Several possibilities for particle control have been proposed to determine which would be most effective in a neurofeedback exercise. This study evaluated these different control options at several public events to determine which artistic representation would be most appropriate for neurofeedback. To do this, we evaluated the artistic aspect of the exercise but also the sensation of particle control and the pleasure of performing the task, which are major motivational parameters to be taken into account in neurofeedback.

## 2. Methods

### 2.1. Population

107 participants, 51 men and 56 women aged 27.6 (±17.1) years on average, participated in the study. Prior to the experiment, oral informed consent was obtained from all participants. The study took place either in laboratory conditions or at various scientific and artistic events of 2018, namely Brain Week, European Researchers’ Night, the VIVO exhibition ‘Entrez en nature !’ and the Hacking Health Besancon. According to French law, this study was classified as a psychology observational study outside of the Jardé law and did not require submission to an ethics committee.

### 2.2. Course of the experiment

At the beginning of the experiment, participants were comfortably seated in a chair, informed of the experimental procedure, and instructed to remain as calm as possible and not to move for the duration of the experiment of about 15 minutes. An EEG headset was then installed with an impedance check lasting about 5 minutes. A 2-minute rest recording was then made to establish a baseline of valence and arousal values. Then, subjects were instructed to try to reach a specific emotional state. Four types of emotional states could be asked of participants, according to Russell’s model (Russell 1980): either a positive valence and a high arousal (emotion of joy or excitement), a positive valence and a low arousal (emotion of calm, relaxation), a negative valence and a high arousal (emotion of irritation, anger), or a negative valence and low arousal (emotion of sadness, fatigue).

### 2.3. Brain data acquisition

The EEG data were acquired from an EEG Emotiv EPOC+ system. This system consists of 16 saline based electrodes and a wireless amplifier. The electrodes are located at positions AF3, F7, F3, FC5, T7, P7, O1, O2, P8, T8, FC6, F4, F8, AF4, according to the international 10-20 system. Two electrodes located just above the ears (P3, P4) are used as a reference. The data is collected at a sampling rate of 128 Hz and transmitted to the computer via Bluetooth.

Although EEG Emotiv systems, which are relatively inexpensive, provide a lower quality signal than when the signal is obtained on more expensive EEG devices (but see (Dikker et al. 2017)), the choice of this material was based on the pragmatic advantages of such a device. The installation time of each Emotiv Epoc+ system is considerably shorter, about 5 minutes, than for gel-based systems, where the gel application for each electrode can ultimately last up to one hour, which considerably extends the duration of the experiments. In addition, since the focus of this study was on evaluating the graphical interface, signal quality was not the main measurement criterion.

### 2.4. Processing of EEG data

EEG processing of valence and arousal is based on methods already used in previous studies (Ramirez et al. 2015; Ramirez et Vamvakousis 2012) using a two-dimensional arousal-valence design (Russell 1980). Data were collected every two seconds. To determine the valence level, the activation levels of the cortical hemispheres were compared. The F3 and F4 electrodes were used to compare alpha activity on the right and left hemispheres because they are located above the prefrontal lobe. Valence was thus calculated by comparing the alpha power at the electrodes F3 and F4, i. e. by applying the following formula: AlphaF4 - AlphaF3. The arousal level was determined by calculating the ratio of beta (12-28 Hz) and alpha (8-12 Hz) oscillations, which may be a reasonable indicator of an individual’s arousal level (Ramirez et al., 2015). The EEG signal was measured on the four electrodes AF3, AF4, F3, F4, which are located above the prefrontal cortex and arousal was calculated as follows: (BetaF3 + BetaF4 + BetaAF3 + BetaAF4) / (AlphaF3 + AlphaF4 + AlphaAF3 + AlphaAF4).

No method of correcting or removing artifacts was applied to the EEG signal. To minimize eye movements, participants were asked to fix the center of the screen during each experiment. To minimize muscle artifacts, participants were asked not to move. If signal quality was not central to this study, in the next steps of performance measurement these parameters will have to be monitored.

### 2.5. Artistic representation of participants’ emotional state

Three types of visual representations were evaluated by participants (Figure 1). All were based on the same principle, namely a representation of emotions in the form of particles tinted according to their location and moving in a white sphere. These particles appeared gradually throughout the experiment.

**Figure 1:**
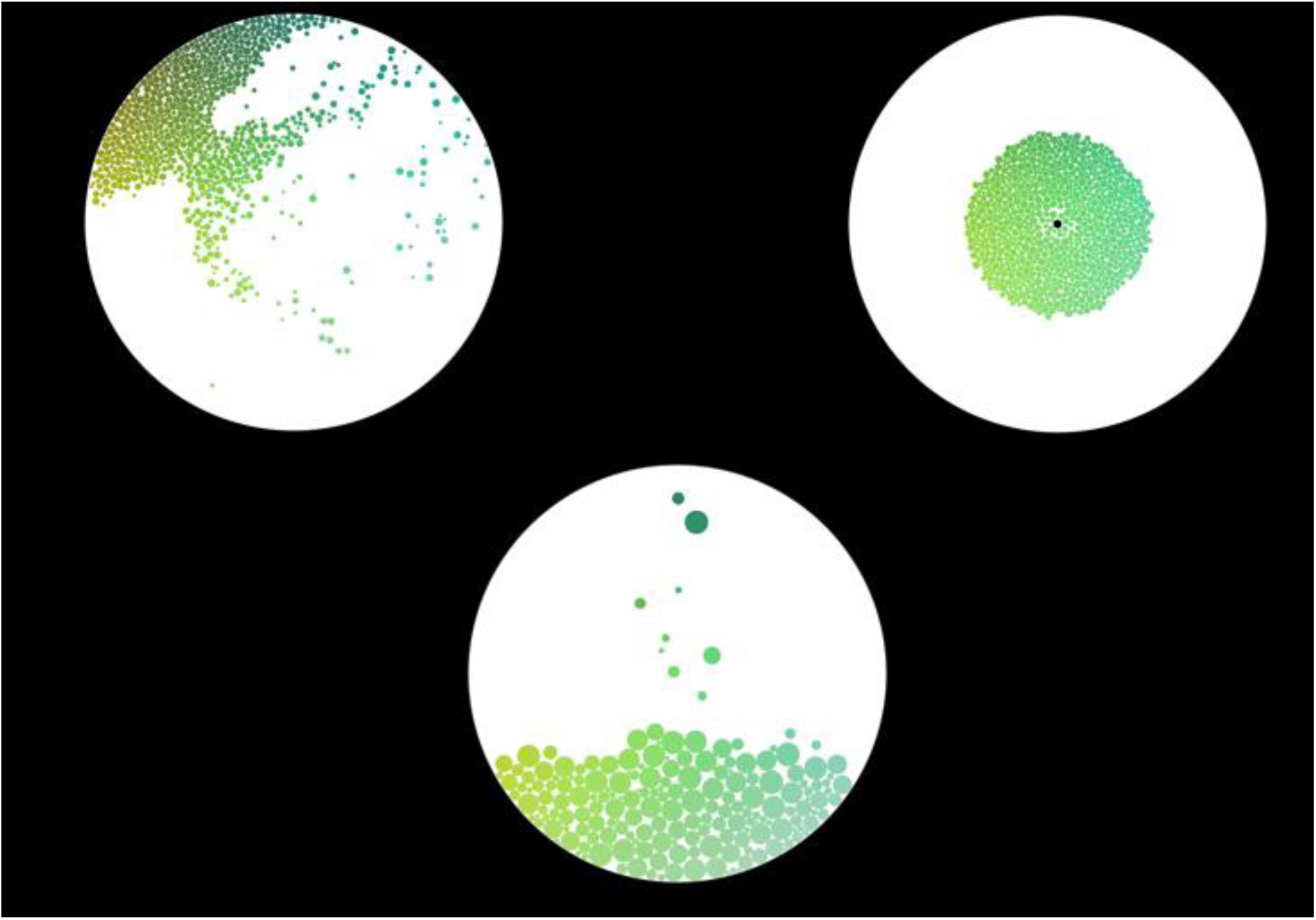
The different types of artistic representation used. In the first representation at the top left, participants had to move the particles to a specific part of the screen. In the second representation at the top right, the participants had to be able to concentrate the particles in the center of the screen. In the third representation at the bottom, the objective was to drop the particles as quickly as possible from the top to the bottom of the screen.

From this common basis, each representation had its own specificities. Each interface played with these particles by modulating the forces applied to them. In the first representation, group of particles moved up or down according to the arousal level, and to the right or left according to the valence level. Thus, for a negative emotion with low arousal, the particles moved to the lower left level of the sphere. Depending on the emotion they were requested to reach, the participants tried to move the particles to a specific part of the sphere. Participants visualized in real time the moving particles in order to give them the feeling of absolute control over their brain activity (see https://youtu.be/c_6IfurxzLc for a video). In the second representation, the objective was to gather the particles in the center of the screen. If the subject was able to modulate his brain activity to the right emotional state, the particles moved towards the center and remained in this position. If the brain activity did not correspond to the requested emotional state then the particles would move back to the periphery (see https://youtu.be/ZeXl43Z7DRU for a video). For the third representation, the objective was to achieve the fastest particle drop from the top to the bottom of the screen until it stuck to the bottom. The more the subject was able to reach the correct emotional state, the faster the particles fell from the top to the bottom of the screen. The more the brain activity moved away from the requested state, the more the particles fell slowly (see https://youtu.be/fuxBEpWwpFA for a video).

The programming of this software is based on the Processing language and intensively uses the physical simulation library adapted for the language by Daniel Shiffman Box2D. The communication between this program and the software that receives and processes the EEG information is done via the OSC protocol, with values ranging between 0 and 100 for valence and arousal. Each visual representation was projected on a circular screen via a video projector. The duration of each exercise was 3 minutes. 89% of the subjects did the same exercise twice, each time with different emotional states to achieve.

### 2.6. Data collection

EEG data collected in real-time were automatically processed to indicate whether the subject had achieved the right emotion during the experiment. For each experiment, the percentage of times a subject had reached the correct level of valence and the correct level of arousal was reported.

In addition, at the end of the experiment, questionnaires were distributed to participants who were asked to indicate on scales between 0 and 5 how the task was artistic, could be used as a clinical activity, whether they felt they had succeeded in controlling particles during the neurofeedback exercise, and whether they had enjoyed the experience.

## 3. Results

### 3.1. Evaluation of each visual representation

45 subjects were tested with visual representation 1, 34 with representation 2 and 28 with representation 3. For one of the users of representation 3, the questionnaire was not completed. The average scores given for each experiment are presented in Figure 2.

**Figure 2:**
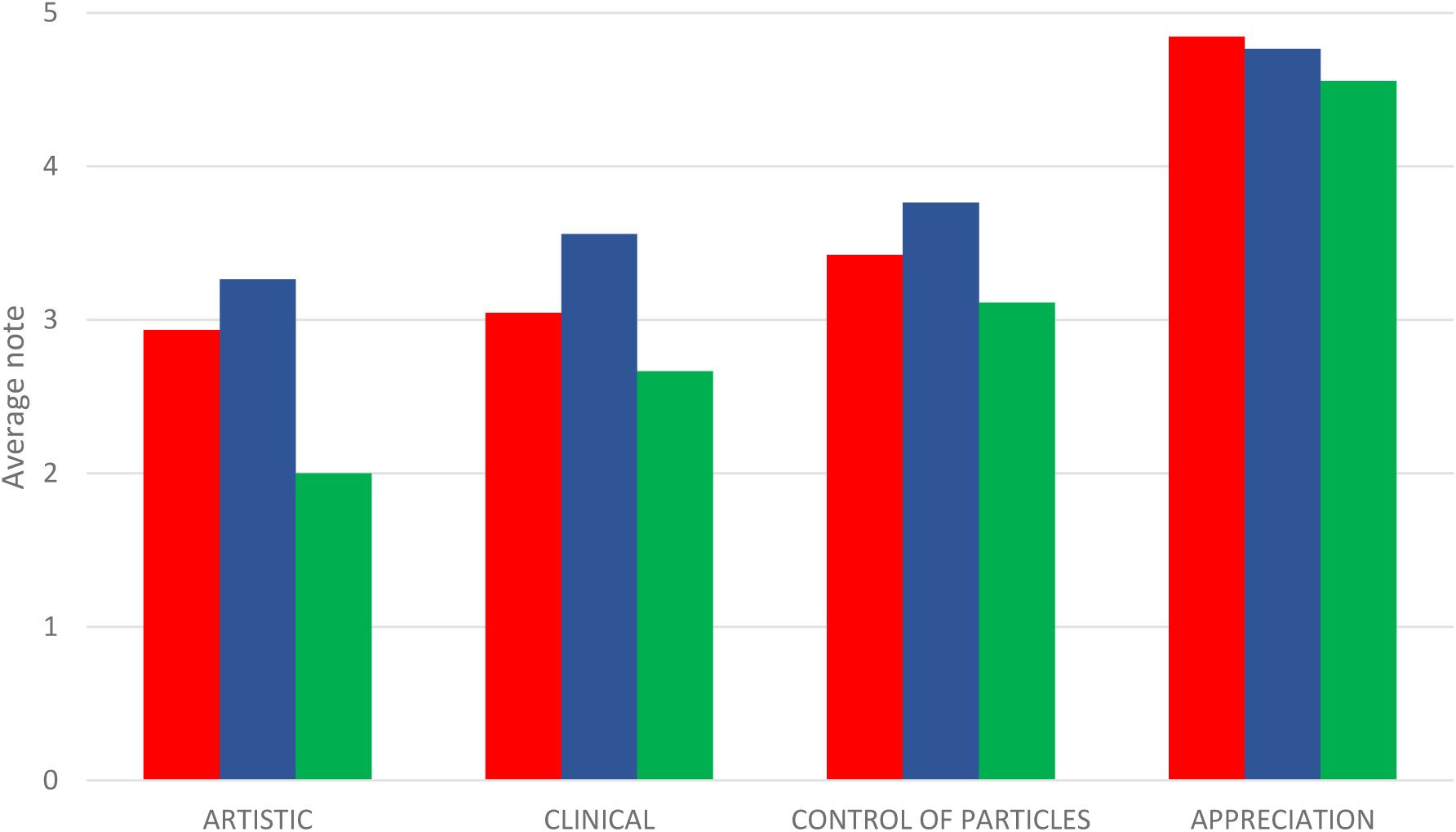
Notes given by the participants during the end-of-experiment questionnaire for each of the questions asked (red: experiment 1, blue: experiment 2, green: experiment 3).

To determine the perception of the audience for the three visual representations, we used non-parametric statistical analyses. With regard to the artistic aspect of the neurofeedback experience, large differences in assessment were observed (Kruskal-Wallis, p=0.0002), the artistic assessment of representation 3 being significantly lower than the other two representations (Mann -Whitney, p=0.004 compared to task 1 and p=0.0002 compared to task 2). For the evaluation of the clinical aspect of each representation, a significant difference was also observed (Kruskal-Wallis, p=0.03). The clinical evaluation of representation 3 was lower than that of representation 2 (Mann -Whitney; p=0.03). To assess whether subjects felt they were in control of the task, only a tendency was observed (Kruskal-Wallis; p=0.06). Finally, with regard to the assessment of the task, very high scores were reported for the 3 tasks, with no significant differences between them (Kruskal-Wallis; p=0.12).

To further explore the difference of artistic perception between the three types of representation, we reported an experience as artistic when participants gave a note of 4 or 5, and non-artistic when participants gave a note of 0 or 1, a method already used before (Zhang et al. 2019). We found that in experience 2, 47% of participants reported having an artistic experience (and 26% a non-artistic experience), whereas there were only 29% of participants in experience 1 (33% not artistic), and 11% in experiment 3 (75% not artistic) (figure 3).

**Figure 3:**
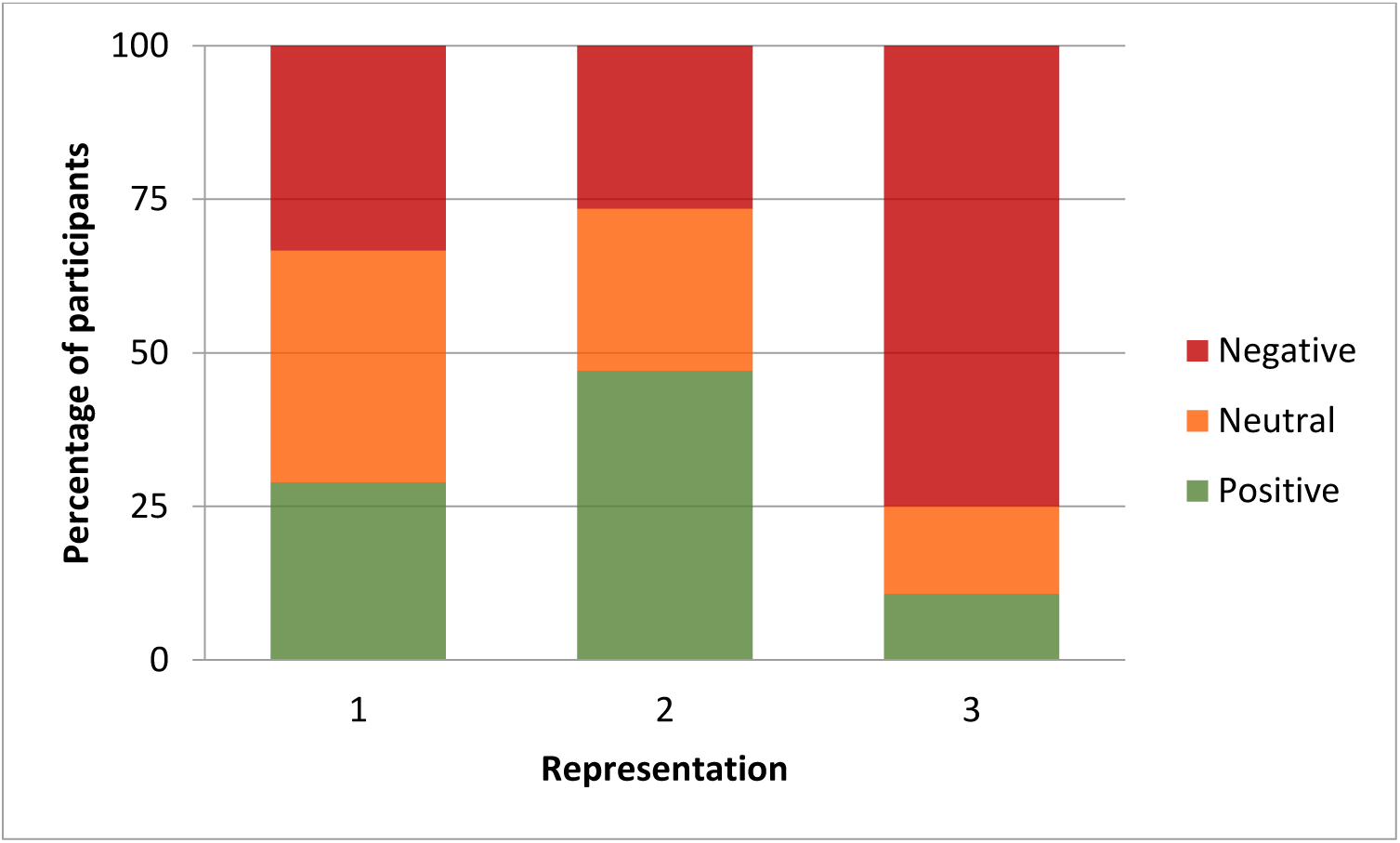
Artistic evaluation of the 3 types of representation. A positive experience was considered as such when participants gave a score of 4 or 5, a partially artistic experience (Neutral) with a score of 3 and a non-artistic (Negative) experience with a score of 1 or 2. In Experiment 2, almost half of the participants reported performing an artistic neurofeedback task.

### 3.2. Neurofeedback evaluation

Although the main purpose of this study was to evaluate the artistic aspect of the device, we did evaluate the participants’ ability to correctly modulate their brain activity during their first session. We compared the percentage of subjects who managed to move the particles according to instruction requested by the experimenter (example: positive valence and positive arousal), and compared the results to the chance level set at 25% (1 in 4 chance of being in the right area). For none of the 3 representations, the subjects were able to significantly reach the correct region (t-test, p>0.1 for all representations).

In the absence of an overall effect, we measured whether subjects were able to control one of the two components (valence or arousal). To study if the subjects had managed to go more easily in one of the components, the chance level was then set at 50%. For valence alone, no representation gave significant results (t-test, p>0.1 for representations 1 and 2; p>0.5 for representation 3). For arousal alone, only the performances of representation 1 were significantly higher than random (t(44)=2.92, p<0.01).

A description of the performance of the subjects for each representation is given in table 1.

**Table 1:** Percentage of participants who were able to place their emotions in the correct area. For the valence and arousal together, the chance level was set at 25%. For the valence alone and for the arousal alone, the chance level was set at 50%. A star means that performance is significantly different from chance at the threshold p<0.05.

|  | valence and<br>arousal | valence alone | arousal alone |
| --- | --- | --- | --- |
| **Experiment 1** | 28,89 | 51,11 | 66,67 * |
| **Experiment 2** | 35,29 | 50,00 | 64,71 |
| **Experiment 3** | 21,43 | 28,57 | 67,86 |

Finally, we compared whether performance improved in subjects who performed the experiment twice (40 subjects for representation 1, 29 for representation 2 and 26 for representation 3). As well overall as for each of the representations separately, no improvement was observed (paired t –test, p>0.1 for all).

## 4. Discussion

The objective of this pilot study was to evaluate whether neurofeedback experiments used in therapeutics could also be artistic. The need for an artistic neurofeedback interface is emerging as the scientific community’s opinion about the effectiveness of neurofeedback is very widely divided (J.-A. Micoulaud-Franchi et al. 2015; Jean-Arthur Micoulaud-Franchi et Fovet 2016; Martijn Arns, Heinrich, et Strehl 2014), with an optimistic part thinking that neurofeedback can be effective and a skeptical part for whom neurofeedback training has no scientific or therapeutic value. Thus it was important that the proposed activity not be seen as just a game and that it really helps to regulate emotions. During the realization of the interface, artists were requested to create a representation, concrete or abstract, of emotions and not only a game unrelated to the final goal of the project (treatment of emotional disorders). Studies have shown that the more playful an application of neurofeedback is, the better the performance (Oude Bos et Reuderink 2008; Bayliss, Inverso, et Tentler 2004), provided that the playful aspect is related to the pathology considered (M. Arns et al. 2017). According to Bandura, people live in a psychological environment that they have largely created themselves (Bandura 1999). Many people are in distress because they are ruminating and cannot control disturbing thoughts. Controlling mental processes is thus a key factor in the self-regulation of emotional states. If neurofeedback is intended to be beneficial to patients by helping them control their own mental processes and therefore their emotional states, having a visual representation of these states can be particularly useful (D. E. J. Linden 2014).

Our goal was to evaluate the most appropriate type of visual feedback to represent emotions. The choice to represent emotions under the form of particles was dictated by the dynamism of this representation. The different forces at play in the organization of particles make them constantly in motion. It is this permanent movement that induces the most interesting aspect; it is the non-punitive response to the objective. The result of the patient’s attempt is not right or wrong, it tends towards or away from the objective, in a natural movement, which can be reminiscent of a lens on a body of water. Particles react with each other to collision, friction, rebound, and sometimes gravity. Of the three representations that were tested, the first two obtained a similar clinical and artistic evaluation, with higher scores for representation 2. In future experiments, the choice of the artistic representation will depend on the purpose of the neurofeedback task: experiment 1 allows visualizing precisely where the subject’s emotional state is located, while experiment 2 mainly gives a binary response (particles are in the center for good emotional state and in the periphery otherwise). Representation 1 requires the integration of the two parameters but has the advantage of being superimposable with the Russell circumplex, which is interesting from a didactic point of view. However, despite similar scores with experiment 1, almost 50% of participants perceived experience 2 as artistic, compared to 30% of participants in experiment 1. Task 2 may better meet the objective of the study, thanks to a visual representation that is less punitive than in other tasks in the event of an error. Therefore, this task may be considered as the entry point in this set of 3 experiments in a therapy framework. Performance 3 resulted in poorer artistic and clinical evaluations, possibly because participants felt a competitive aspect in having to drop the balls quickly. In addition, in this representation, it is more difficult to know if brain activity is well controlled since the participant does not have a particle fall rate reference on which to refer. It is interesting to note how the modification of a single parameter, in this case the gravitational force, can have major implications on the artistic and clinical perception of the task. In terms of the pleasure of completing the task, the results of the three representations are encouraging since the subjects greatly appreciated participating in the experiment and had the impression that they were controlling the particles. These parameters are important to motivate a subject to repeat a neurofeedback experiment, because in some cases, more than 30 sessions are required to demonstrate neurofeedback effectiveness (Marzbani, Marateb, et Mansourian 2016).

Artistic scores may not be considered as very high, with the average score in experiments 1 and 2 being around 3/5, but it is important to note that the aim was not to obtain the highest possible artistic score, but to achieve a task that is perceived as artistic and applicable in clinical practice. With the development of portable and relatively inexpensive EEG systems, there are now a number of projects that have associated art with sensory feedback, with no real possibility of clinical application. For example, the GlobalMind project sought to combine art and EEG activity to generate audiovisual effects. This has led to the production of ‘Spectacle of the mind’ shows presented to the general public. Another example is the Ascent project (www.theascent.co) where an installation allows individuals to levitate by modulating their ability to concentrate around an auditory and luminous show. It is by controlling this activity that participants can climb more or less high in the air. In addition to these exclusively artistic uses of real time feedback in EEG, other approaches have been used with both an artistic and pedagogical focus. For example, the project “My Virtual Dream” which was presented in Toronto in 2013, at the Nuits Blanches art festival, and which measured EEG activities of 523 participants in a single night (Kovacevic et al. 2015). Participants practiced simple EEG tasks targeting either a state of relaxation or a state of concentration. During the evening, an improvement in performance was observed, observable after only 1 minute of training. A dome that allows spatialization of the individual’s brain activities and has also been developed to improve the individual’s immersive appearance compared to a simple screen (Grandchamp et Delorme 2016). This tool is intended to illustrate scientific knowledge about the brain. The authors also believe that this type of artistic and immersive environment would increase patients’ motivation while reducing their training time and fatigue. Of course such a dome remains difficult to use in common clinical practice.

In this study it is important to dissociate the emotion measurement device from the representation interface. As the main purpose of this neurofeedback pilot was to evaluate the artistic aspect of the interface, data processing, signal filtering, real-time rejection of flashes and eye or muscle movements were not optimally exploited, although they can have a major influence on the quality of EEG plots. Similarly, it is possible to improve the quality of the EEG signal by using gel-based electrodes. Here we measured emotions with the material and parameters already described in the literature to calculate valence and arousal (Ramirez et al. 2015). However, it is important to note that this interface could be used with different methods of measuring emotions and on different populations. For example, there are other methods for detecting emotions in EEGs, for example with connectivity analysis (Koush et al. 2017). In this case, subjects must regulate the top-down activity of the prefrontal cortex to the amygdala. This artistic interface could also be applied to fMRI, the other major neuroimaging method for measuring emotions. Although the fMRI technique provides only indirect measurements of neural activity and has a much lower temporal resolution than the EEG, its spatial resolution and access to deeper structures make it an attractive tool for network mapping and neurofeedback. Depending on the method chosen and the brain region targeted, this emotional measurement interface could potentially be applied for the treatment of mental disorders such as depression, schizophrenia or bipolar disorders.

This study is the first step and several points remain to be clarified to test the effectiveness of this type of artistic representation. First, it is not known how well the subjects were trying to achieve the requested emotional state. We did not control the extent to which subjects used different strategies among themselves and over time, which can strongly influence neurofeedback performance. Moreover, from a methodological point of view, it will be necessary to establish a control condition, a critical point in any neurofeedback study, to verify whether the effect comes from the experience itself, or from other factors such as the attention given to the patient (Thibault et Raz 2017; Jean-Arthur Micoulaud-Franchi et Fovet 2018). In addition, it is well known that the placebo effect can have a significant influence on the outcome. However, if the result is present, the use of such a method may be acceptable, even as a placebo (Thibault et Raz 2016). Finally, future evaluations will have to assess whether the artistic interface manages to keep the level of motivation of participants at a high level during repeated experiences. Although all subjects strongly appreciated performing the experiment, and felt that they were controlling the particles, it is likely that this motivation will gradually decrease and will need to be assessed in comparison to other types of visual feedback.

## Conclusion

In this pilot study involving the collaboration between neuroscientists and digital artists, we were able to set up a neurofeedback interface for emotion regulation that is perceived as both an artistic and clinical activity. It will remain to be explored whether the therapeutic effect of neurofeedback can make clinical sense and how to carry out a neurofeedback examination in an optimal way. For this reason, the design of appropriate control conditions for clinical trials is a real challenge. It will also be necessary to identify precisely the patient populations for which neurofeedback can work. The cognitive and motivational factors underlying effective neurofeedback training are largely unknown. For example, if this interface is to be applied to patients suffering from anhedonia, the sub-components causing the anhedonic disorder should be well separated because they may originate in different brain regions (Thomsen 2015).

## References

Aftanas, L. I., et S. A. Golocheikine. 2001. « Human Anterior and Frontal Midline Theta and Lower Alpha Reflect Emotionally Positive State and Internalized Attention: High-Resolution EEG Investigation of Meditation ». Neuroscience Letters 310 (1): 57–60. https://doi.org/10.1016/s0304-3940(01)02094-8.

Arjmand, Hussain-Abdulah, Jesper Hohagen, Bryan Paton, et Nikki S. Rickard. 2017. « Emotional Responses to Music: Shifts in Frontal Brain Asymmetry Mark Periods of Musical Change ». Frontiers in Psychology 8: 2044. https://doi.org/10.3389/fpsyg.2017.02044.

Arns, M., J.-M. Batail, S. Bioulac, M. Congedo, C. Daudet, D. Drapier, T. Fovet, et al. 2017. « Neurofeedback: One of Today’s Techniques in Psychiatry? » L’Encephale 43 (2): 135–45. https://doi.org/10.1016/j.encep.2016.11.003.

Arns, Martijn, Hartmut Heinrich, et Ute Strehl. 2014. « Evaluation of Neurofeedback in ADHD: The Long and Winding Road ». Biological Psychology 95 (janvier): 108–15. https://doi.org/10.1016/j.biopsycho.2013.11.013.

Baehr, E., et R Baehr. 1997. « The Use of Brainwave Biofeedback as an Adjunctive Therapeutic Treatment for Depression: Three Case Studies ». Biofeedback 25 (1): 10–11.

Baehr, E., J.P. Rosenfeld, et R Baehr. 1997. «. The clinical use of an alpha asymmetry protocol in the neurofeedback treatment of depression: two case studies ». J. Neurotherapy 2: 10–23.

Bagdasaryan, Juliana, et Michel Le Van Quyen. 2013. « Experiencing Your Brain: Neurofeedback as a New Bridge between Neuroscience and Phenomenology ». Frontiers in Human Neuroscience 7: 680. https://doi.org/10.3389/fnhum.2013.00680.

Bandura, A. 1999. « Moral Disengagement in the Perpetration of Inhumanities ». Personality and Social Psychology Review: An Official Journal of the Society for Personality and Social Psychology, Inc 3 (3): 193–209. https://doi.org/10.1207/s15327957pspr0303_3.

Bayliss, JD, SA Inverso, et A Tentler. 2004. « Changing the P300 brain computer interface ». CyberPsychol Behav 7 (6): 694–704.

Caria, Andrea, Ranganatha Sitaram, Ralf Veit, Chiara Begliomini, et Niels Birbaumer. 2010. « Volitional Control of Anterior Insula Activity Modulates the Response to Aversive Stimuli. A Real-Time Functional Magnetic Resonance Imaging Study ». Biological Psychiatry 68 (5): 425–32. https://doi.org/10.1016/j.biopsych.2010.04.020.

Caria, Andrea, Ralf Veit, Ranganatha Sitaram, Martin Lotze, Nikolaus Weiskopf, Wolfgang Grodd, et Niels Birbaumer. 2007. « Regulation of Anterior Insular Cortex Activity Using Real-Time FMRI ». NeuroImage 35 (3): 1238–46. https://doi.org/10.1016/j.neuroimage.2007.01.018.

Cavazza, Marc, G Aranyi, Fred Charles, Julie Porteous, Stephen Gilroy, Ilana Klovatch, Gilan Jackont, et al. 2014. « Towards empathic neurofeedback for interactive storytelling ». Open Access Series in Informatics, 42–60.

Choi, Sung Won, Sang Eun Chi, Sun Yong Chung, Jong Woo Kim, Chang Yil Ahn, et Hyun Taek Kim. 2011. « Is Alpha Wave Neurofeedback Effective with Randomized Clinical Trials in Depression? A Pilot Study ». Neuropsychobiology 63 (1): 43–51. https://doi.org/10.1159/000322290.

Coan, James A., et John J. B. Allen. 2004. « Frontal EEG Asymmetry as a Moderator and Mediator of Emotion ». Biological Psychology 67 (1-2): 7–49. https://doi.org/10.1016/j.biopsycho.2004.03.002.

Cook, I. A., R. O’Hara, S. H. Uijtdehaage, M. Mandelkern, et A. F. Leuchter. 1998. « Assessing the Accuracy of Topographic EEG Mapping for Determining Local Brain Function ». Electroencephalography and Clinical Neurophysiology 107 (6): 408–14. https://doi.org/10.1016/s0013-4694(98)00092-3.

Cunningham, William A., Nathan L. Arbuckle, Andrew Jahn, Samantha M. Mowrer, et Amir M. Abduljalil. 2010. « Aspects of Neuroticism and the Amygdala: Chronic Tuning from Motivational Styles ». Neuropsychologia 48 (12): 3399–3404. https://doi.org/10.1016/j.neuropsychologia.2010.06.026.

Cunningham, William A., Carol L. Raye, et Marcia K. Johnson. 2005. « Neural Correlates of Evaluation Associated with Promotion and Prevention Regulatory Focus ». Cognitive, Affective & Behavioral Neuroscience 5 (2): 202–11.

Davidson, R. J. 1988. « EEG Measures of Cerebral Asymmetry: Conceptual and Methodological Issues ». The International Journal of Neuroscience 39 (1-2): 71–89. https://doi.org/10.3109/00207458808985694.

Davidson, R. J. 1992. « Anterior Cerebral Asymmetry and the Nature of Emotion ». Brain and Cognition 20 (1): 125–51.

Davidson, R. J. 1998. « Anterior Electrophysiological Asymmetries, Emotion, and Depression: Conceptual and Methodological Conundrums ». Psychophysiology 35 (5): 607–14. https://doi.org/10.1017/s0048577298000134.

Davidson, R. J., P. Ekman, C. D. Saron, J. A. Senulis, et W. V. Friesen. 1990. « Approach-Withdrawal and Cerebral Asymmetry: Emotional Expression and Brain Physiology. I ». Journal of Personality and Social Psychology 58 (2): 330–41.

deCharms, R. Christopher. 2008. « Applications of Real-Time FMRI ». Nature Reviews. Neuroscience 9 (9): 720–29. https://doi.org/10.1038/nrn2414.

Dikker, Suzanne, Lu Wan, Ido Davidesco, Lisa Kaggen, Matthias Oostrik, James McClintock, Jess Rowland, et al. 2017. « Brain-to-Brain Synchrony Tracks Real-World Dynamic Group Interactions in the Classroom ». Current Biology: CB 27 (9): 1375–80. https://doi.org/10.1016/j.cub.2017.04.002.

Ertl, Matthias, Maria Hildebrandt, Kristina Ourina, Gregor Leicht, et Christoph Mulert. 2013. « Emotion Regulation by Cognitive Reappraisal - the Role of Frontal Theta Oscillations ». NeuroImage 81 (novembre): 412–21. https://doi.org/10.1016/j.neuroimage.2013.05.044.

Gaume, A., A. Vialatte, A. Mora-Sánchez, C. Ramdani, et F. B. Vialatte. 2016. « A Psychoengineering Paradigm for the Neurocognitive Mechanisms of Biofeedback and Neurofeedback ». Neuroscience and Biobehavioral Reviews 68 (septembre): 891–910. https://doi.org/10.1016/j.neubiorev.2016.06.012.

Grandchamp, Romain, et Arnaud Delorme. 2016. « The Brainarium: An Interactive Immersive Tool for Brain Education, Art, and Neurotherapy ». Computational Intelligence and Neuroscience 2016: 4204385. https://doi.org/10.1155/2016/4204385.

Gruzelier, John H. 2014. « EEG-Neurofeedback for Optimising Performance. I: A Review of Cognitive and Affective Outcome in Healthy Participants ». Neuroscience and Biobehavioral Reviews 44 (juillet): 124–41. https://doi.org/10.1016/j.neubiorev.2013.09.015.

Gruzelier, John H. 2018. « Enhancing Creativity with Neurofeedback in the Performing Arts: Actors, Musicians, Dancers. » In Burgoyne S. (eds) Creativity in Theatre. Creativity Theory and Action in Education. Vol. 2. Springer, Cham.

Harmon-Jones, Eddie, et Philip A. Gable. 2018. « On the Role of Asymmetric Frontal Cortical Activity in Approach and Withdrawal Motivation: An Updated Review of the Evidence ». Psychophysiology 55 (1). https://doi.org/10.1111/psyp.12879.

Kinreich, Sivan, Ilana Podlipsky, Shahar Jamshy, Nathan Intrator, et Talma Hendler. 2014. « Neural Dynamics Necessary and Sufficient for Transition into Pre-Sleep Induced by EEG Neurofeedback ». NeuroImage 97 (août): 19–28. https://doi.org/10.1016/j.neuroimage.2014.04.044.

Koush, Yury, Djalel-E. Meskaldji, Swann Pichon, Gwladys Rey, Sebastian W. Rieger, David E. J. Linden, Dimitri Van De Ville, Patrik Vuilleumier, et Frank Scharnowski. 2017. « Learning Control Over Emotion Networks Through Connectivity-Based Neurofeedback ». Cerebral Cortex (New York, N.Y.: 1991) 27 (2): 1193–1202. https://doi.org/10.1093/cercor/bhv311.

Kovacevic, Natasha, Petra Ritter, William Tays, Sylvain Moreno, et Anthony Randal McIntosh. 2015. « “My Virtual Dream”: Collective Neurofeedback in an Immersive Art Environment ». PloS One 10 (7): e0130129. https://doi.org/10.1371/journal.pone.0130129.

Linden, David. 2013. « Biological Psychiatry: Time for New Paradigms ». The British Journal of Psychiatry: The Journal of Mental Science 202 (3): 166–67. https://doi.org/10.1192/bjp.bp.112.121269.

Linden, David E. J. 2014. « Neurofeedback and Networks of Depression ». Dialogues in Clinical Neuroscience 16 (1): 103–12.

Lorenzetti, Valentina, Bruno Melo, Rodrigo Basílio, Chao Suo, Murat Yücel, Carlos J. Tierra-Criollo, et Jorge Moll. 2018. « Emotion Regulation Using Virtual Environments and Real-Time FMRI Neurofeedback ». Frontiers in Neurology 9: 390. https://doi.org/10.3389/fneur.2018.00390.

Marzbani, Hengameh, Hamid Reza Marateb, et Marjan Mansourian. 2016. « Neurofeedback: A Comprehensive Review on System Design, Methodology and Clinical Applications ». Basic and Clinical Neuroscience 7 (2): 143–58. https://doi.org/10.15412/J.BCN.03070208.

Meir-Hasson, Yehudit, Sivan Kinreich, Ilana Podlipsky, Talma Hendler, et Nathan Intrator. 2014. « An EEG Finger-Print of FMRI Deep Regional Activation ». NeuroImage 102 Pt 1 (novembre): 128–41. https://doi.org/10.1016/j.neuroimage.2013.11.004.

Micoulaud-Franchi, J.-A., A. McGonigal, R. Lopez, C. Daudet, I. Kotwas, et F. Bartolomei. 2015. « Electroencephalographic Neurofeedback: Level of Evidence in Mental and Brain Disorders and Suggestions for Good Clinical Practice ». Neurophysiologie Clinique = Clinical Neurophysiology 45 (6): 423–33. https://doi.org/10.1016/j.neucli.2015.10.077.

Micoulaud-Franchi, Jean-Arthur, et Thomas Fovet. 2016. « Neurofeedback: Time Needed for a Promising Non-Pharmacological Therapeutic Method ». The Lancet. Psychiatry 3 (9): e16. https://doi.org/10.1016/S2215-0366(16)30189-4.

Micoulaud-Franchi, Jean-Arthur, et Thomas Fovet. 2018. « A Framework for Disentangling the Hyperbolic Truth of Neurofeedback: Comment on Thibault and Raz (2017) ». The American Psychologist 73 (7): 933–35. https://doi.org/10.1037/amp0000340.

Oude Bos, D. et B Reuderink. 2008. « BrainBasher: A BCI game. » In Extended Abstracts of the International Conference on Fun and Games 2008, Eindhoven, Netherlands, 36–39. Eindhoven University of Technology, Eindhoven, The Netherlands.

Papousek, Ilona, Elisabeth M. Weiss, Günter Schulter, Andreas Fink, Eva M. Reiser, et Helmut K. Lackner. 2014. « Prefrontal EEG Alpha Asymmetry Changes While Observing Disaster Happening to Other People: Cardiac Correlates and Prediction of Emotional Impact ». Biological Psychology 103 (décembre): 184–94. https://doi.org/10.1016/j.biopsycho.2014.09.001.

Paquette, Vincent, Mario Beauregard, et Dominic Beaulieu-Prévost. 2009. « Effect of a Psychoneurotherapy on Brain Electromagnetic Tomography in Individuals with Major Depressive Disorder ». Psychiatry Research 174 (3): 231–39. https://doi.org/10.1016/j.pscychresns.2009.06.002.

Peeters, Frenk, Mare Oehlen, Jacco Ronner, Jim van Os, et Richel Lousberg. 2014. « Neurofeedback as a Treatment for Major Depressive Disorder--a Pilot Study ». PloS One 9 (3): e91837. https://doi.org/10.1371/journal.pone.0091837.

Pizzagalli, Diego A., Jack B. Nitschke, Terrence R. Oakes, Andrew M. Hendrick, Kathryn A. Horras, Christine L. Larson, Heather C. Abercrombie, et al. 2002. « Brain Electrical Tomography in Depression: The Importance of Symptom Severity, Anxiety, and Melancholic Features ». Biological Psychiatry 52 (2): 73–85. https://doi.org/10.1016/s0006-3223(02)01313-6.

Ramirez, Rafael, Manel Palencia-Lefler, Sergio Giraldo, et Zacharias Vamvakousis. 2015. « Musical Neurofeedback for Treating Depression in Elderly People ». Frontiers in Neuroscience 9: 354. https://doi.org/10.3389/fnins.2015.00354.

Ramirez, Rafael, et Zacharias Vamvakousis. 2012. « Detecting emotion from EEG signals using the emotive epoc device ». In Proceedings of the 2012 International Conference on Brain Informatics, LNCS 7670, 175–84. Macau: Springer.

Ruiz, Sergio, Sangkyun Lee, Surjo R. Soekadar, Andrea Caria, Ralf Veit, Tilo Kircher, Niels Birbaumer, et Ranganatha Sitaram. 2013. « Acquired Self-Control of Insula Cortex Modulates Emotion Recognition and Brain Network Connectivity in Schizophrenia ». Human Brain Mapping 34 (1): 200–212. https://doi.org/10.1002/hbm.21427.

Russell, James A. 1980. « A circumplex model of affect ». Journal of Personality and Social Psychology 39: 1161–78.

Schlund, Michael W., et Michael F. Cataldo. 2010. « Amygdala Involvement in Human Avoidance, Escape and Approach Behavior ». NeuroImage 53 (2): 769–76. https://doi.org/10.1016/j.neuroimage.2010.06.058.

Shtark, M. B., E. G. Verevkin, L. I. Kozlova, K. G. Mazhirina, M. A. Pokrovskii, E. D. Petrovskii, A. A. Savelov, A. S. Starostin, et S. V. Yarosh. 2015. « Synergetic FMRI-EEG Brain Mapping in Alpha-Rhythm Voluntary Control Mode ». Bulletin of Experimental Biology and Medicine 158 (5): 644–49. https://doi.org/10.1007/s10517-015-2827-7.

Spielberg, Jeffrey M., Gregory A. Miller, Stacie L. Warren, Anna S. Engels, Laura D. Crocker, Marie T. Banich, Bradley P. Sutton, et Wendy Heller. 2012. « A Brain Network Instantiating Approach and Avoidance Motivation ». Psychophysiology 49 (9): 1200–1214. https://doi.org/10.1111/j.1469-8986.2012.01443.x.

Sulzer, J., S. Haller, F. Scharnowski, N. Weiskopf, N. Birbaumer, M. L. Blefari, A. B. Bruehl, et al. 2013. « Real-Time FMRI Neurofeedback: Progress and Challenges ». NeuroImage 76 (août): 386–99. https://doi.org/10.1016/j.neuroimage.2013.03.033.

Sutton, Steven K., et Richard J. Davidson. 1997. « Prefrontal Brain Asymmetry: A Biological Substrate of the Behavioral Approach and Inhibition Systems ». Psychological Science 8 (3): 204–10. https://doi.org/10.1111/j.1467-9280.1997.tb00413.x.

Thibault, Robert T., et Amir Raz. 2016. « When Can Neurofeedback Join the Clinical Armamentarium? » The Lancet. Psychiatry 3 (6): 497–98. https://doi.org/10.1016/S2215-0366(16)30040-2.

Thibault, Robert T., et Amir Raz. 2017. « The Psychology of Neurofeedback: Clinical Intervention Even If Applied Placebo ». The American Psychologist 72 (7): 679–88. https://doi.org/10.1037/amp0000118.

Thomsen, Kristine Rømer. 2015. « Measuring Anhedonia: Impaired Ability to Pursue, Experience, and Learn about Reward ». Frontiers in Psychology 6: 1409. https://doi.org/10.3389/fpsyg.2015.01409.

Weiskopf, Nikolaus. 2012. « Real-Time FMRI and Its Application to Neurofeedback ». NeuroImage 62 (2): 682–92. https://doi.org/10.1016/j.neuroimage.2011.10.009.

Weiskopf, Nikolaus, Frank Scharnowski, Ralf Veit, Rainer Goebel, Niels Birbaumer, et Klaus Mathiak. 2004. « Self-Regulation of Local Brain Activity Using Real-Time Functional Magnetic Resonance Imaging (FMRI) ». Journal of Physiology, Paris 98 (4-6): 357–73. https://doi.org/10.1016/j.jphysparis.2005.09.019.

Wheeler, R. E., R. J. Davidson, et A. J. Tomarken. 1993. « Frontal Brain Asymmetry and Emotional Reactivity: A Biological Substrate of Affective Style ». Psychophysiology 30 (1): 82–89. https://doi.org/10.1111/j.1469-8986.1993.tb03207.x.

Young, Kymberly D., Vadim Zotev, Raquel Phillips, Masaya Misaki, Han Yuan, Wayne C. Drevets, et Jerzy Bodurka. 2014. « Real-Time FMRI Neurofeedback Training of Amygdala Activity in Patients with Major Depressive Disorder ». PloS One 9 (2): e88785. https://doi.org/10.1371/journal.pone.0088785.

Yuan, Han, Kymberly D. Young, Raquel Phillips, Vadim Zotev, Masaya Misaki, et Jerzy Bodurka. 2014. « Resting-State Functional Connectivity Modulation and Sustained Changes after Real-Time Functional Magnetic Resonance Imaging Neurofeedback Training in Depression ». Brain Connectivity 4 (9): 690–701. https://doi.org/10.1089/brain.2014.0262.

Zhang, Jack, Zeanna Jadavji, Ephrem Zewdie, et Adam Kirton. 2019. « Evaluating If Children Can Use Simple Brain Computer Interfaces ». Frontiers in Human Neuroscience 13: 24. https://doi.org/10.3389/fnhum.2019.00024.

Zich, Catharina, Stefan Debener, Cornelia Kranczioch, Martin G. Bleichner, Ingmar Gutberlet, et Maarten De Vos. 2015. « Real-Time EEG Feedback during Simultaneous EEG-FMRI Identifies the Cortical Signature of Motor Imagery ». NeuroImage 114 (juillet): 438–47. https://doi.org/10.1016/j.neuroimage.2015.04.020.

Zotev, Vadim, Frank Krueger, Raquel Phillips, Ruben P. Alvarez, W. Kyle Simmons, Patrick Bellgowan, Wayne C. Drevets, et Jerzy Bodurka. 2011. « Self-Regulation of Amygdala Activation Using Real-Time FMRI Neurofeedback ». PloS One 6 (9): e24522. https://doi.org/10.1371/journal.pone.0024522.

Zotev, Vadim, Raquel Phillips, Kymberly D. Young, Wayne C. Drevets, et Jerzy Bodurka. 2013. « Prefrontal Control of the Amygdala during Real-Time FMRI Neurofeedback Training of Emotion Regulation ». PloS One 8 (11): e79184. https://doi.org/10.1371/journal.pone.0079184.

Zotev, Vadim, Han Yuan, Masaya Misaki, Raquel Phillips, Kymberly D. Young, Matthew T. Feldner, et Jerzy Bodurka. 2016. « Correlation between Amygdala BOLD Activity and Frontal EEG Asymmetry during Real-Time FMRI Neurofeedback Training in Patients with Depression ». NeuroImage. Clinical 11: 224–38. https://doi.org/10.1016/j.nicl.2016.02.003.

